# Networks of VTA neurons encode real-time information about uncertain numbers of actions executed to earn a reward

**DOI:** 10.1101/084970

**Authors:** Jesse Wood, Nicholas W. Simon, Spencer Koerner, Robert E. Kass, Bita Moghaddam

**Affiliations:** Department of Psychiatry, University of Pittsburgh, 450 Technology Drive Suite 223, Pittsburgh, PA 15219.; Department of Psychology, University of Memphis, Psychology Building 400 Innovation Drive, Memphis, TN 38152; Department of Statistics, Carnegie Mellon University, 132 Baker Hall, Pittsburgh, PA 15213.; Center for the Neural Basis of Cognition, Carnegie Mellon University and the University of Pittsburgh, 4400 5^th^ Ave, Pittsburgh PA, 15213; Machine Learning Department, Carnegie Mellon University, Gates Hillman Center 8203, Pittsburgh, PA 15213; Department of Behavioral Neuroscience, Oregon Health and Sciences University, Portland, OR 97239

## Abstract

Multiple and unpredictable numbers of actions are often required to achieve a goal. In order to organize behavior and allocate effort so that optimal behavioral policies can be selected, it is necessary to continually monitor ongoing actions. Real-time processing of information related to actions and outcomes is typically assigned to the prefrontal cortex and basal ganglia, but also depends on midbrain regions, especially the ventral tegmental area (VTA). We were interested in how individual VTA neurons, as well as networks within the VTA, encode salient events when an unpredictable number of serial actions are required to obtain a reward. We recorded from ensembles of putative dopamine and non-dopamine neurons in the VTA as animals performed multiple cued trials in a recording session where, in each trial, serial actions were randomly rewarded. While averaging population activity did not reveal a response pattern, we observed that different neurons selectively tuned to low, medium, or high numbered actions in a trial. This preferential tuning of putative dopamine and non-dopamine VTA neurons to different subsets of actions in a trial allowed information about binned action number to be decoded from the ensemble activity. At the network level, tuning curve similarity was positively associated with action-evoked noise correlations, suggesting that action number selectivity reflects functional connectivity within these networks. Analysis of phasic responses to cue and reward revealed that the requirement to execute multiple and uncertain numbers of actions weakens both cue-evoked responses and cue-reward response correlation. The functional connectivity and ensemble coding scheme that we observe here may allow VTA neurons to cooperatively provide a real-time account of ongoing behavior. These computations may be critical to cognitive and motivational functions that have long been associated with VTA dopamine neurons.

## INTRODUCTION

Organization of goal-directed behavior requires real-time monitoring of ongoing actions. This function is typically assigned to the prefrontal cortex and striatum, but several lines of evidence demonstrate that organization of a series of behaviors also depends on VTA dopamine input to these regions (Goldman-Rakic, 1988; Robbins and Arnsten, 2009; Salamone et al., 2009). For instance, lesions of VTA dopamine projections to the striatum, or antagonists of dopamine receptors, impair the capacity to perform large numbers of actions without impacting the ability to complete one (or a few) action(s) (Aberman and Salamone, 1999; Ishiwari et al., 2004; Mingote et al., 2005). Additionally, VTA neurons are implicated in several cognitive constructs necessary for organizing an ongoing series of behaviors, such as working memory, attention, effort, and cognitive flexibility (Salamone and Correa, 2002; Seamans and Yang, 2004; Robbins and Arnsten, 2009; Fujisawa and Buzsaki, 2011). Thus, VTA neurons are critical for organizing, sustaining, and adapting an ongoing sequence of behaviors.

Additional evidence that VTA neurons participate in structuring and organizing a behavioral series is provided by studies showing that dopamine release in nucleus accumbens precedes a series of two lever presses (Wassum et al., 2012), and dopamine neurons encode the first and last actions in a fixed response eight (FR8) reinforcement schedule (Jin and Costa, 2010). These response patterns, however, do not reveal how VTA neurons encode information *during* a series of actions. The expansive literature on dopamine signaling has mostly used tasks that require subjects to execute a single instrumental action, or Pavlovian tasks with no explicit action requirement. These experiments have generally focused on neuronal responses before behavior is initiated or after outcomes are delivered (Ljungberg et al., 1992; Schultz et al., 1993; Montague et al., 1996; Schultz et al., 1997; Schultz, 1998; Bayer and Glimcher, 2005; Morris et al., 2006; Roesch et al., 2007; Salamone et al., 2007; Dayan and Niv, 2008; Glimcher, 2011). Thus, little is known about how VTA neurons encode information that could be used to organize and sustain a series of actions.

To begin to understand how VTA neurons monitor a series of actions in real-time, we recorded from VTA ensembles during a task where rats executed an unpredictable number of self-paced actions to earn reward. We found that individual dopamine and non-dopamine neurons fired selectively during different actions in a trial. Whereas population-averaging concealed this mosaic of activity patterns, it was possible to decode low, medium, and high numbered actions within a trial from the collective activity of all neurons. These data provide novel evidence of ensemble encoding by VTA neurons and provide critical insight into how VTA neurons may contribute to complex behaviors by encoding real-time information about ongoing actions.

## MATERIALS AND METHODS

### Subjects and Apparatus

All procedures were conducted in accordance with the National Institutes of Health’s Guide to the Care and Use of Laboratory Animals, and approved by the University of Pittsburgh’s Institutional Animal Care and Use Committee. Behavioral and neuronal data were collected from 10 adult, experimentally naïve, male Sprague-Dawley rats (Harlan, Frederick, MD) singly housed on a 12-hour light cycle (lights on at 7pm). These sample sizes were chosen because they are sufficient to detect behavioral and VTA neuronal effects in a variety of rat operant procedures in our lab (Kim et al., 2010; Kim et al., 2012; Kim et al., 2016). At the time of surgery, rats weighed approximately 350 grams (approximately 12 weeks of age). Under isoflurane anesthesia, 8 or 16 channel, 50 μm stainless steel, Teflon insulated, microelectrode arrays were implanted in left VTA (relative to Bregma: -5.30 mm posterior, 0.8 mm lateral, and 8.3 mm ventral). All experiments were conducted in a standard operant chamber (Coulbourn Instruments, Allentown, PA). The operant chamber had a food trough on one wall and a single nose poke port on the opposite wall. Both the food trough and nose poke port could be illuminated and were equipped with an infrared beam, which detected the animal’s entry. The operant chamber system controller was configured to send the time of behavioral and environmental events to the recording interface via standard TTL pulses and a digital interface.

### Behavior

Each rat was given 7 days to recover from surgery and food restricted to 90% of their free feeding body weights. Rats were habituated to handling for 5 minutes per day for 3 consecutive days, before being habituated to being handled and connected to a headstage cable in the procedure room for 2 additional days. Following habituation, rats were given a single 30 minute magazine training session in the operant chamber, in which sugar pellets were delivered on a variable time 75s reinforcement schedule. When each pellet was delivered, the pellet trough was illuminated for 4s. The animal’s behavior had no programmed consequences in the magazine training session.

Following the magazine training session, each animal began instrumental conditioning. During all instrumental conditioning sessions, each trial began with illumination of the nose poke port (cue light onset). This served as a discriminative stimulus that reinforcing outcomes (sugar pellets) were available (termed the ‘response period’), contingent upon the animal executing actions (nose pokes into the lit port). In each trial, actions were reinforced randomly, according to a predetermined probability. When an action was executed, the behavioral system controller randomly drew an outcome state (either reinforcement or no programmed consequence) with replacement, according to the probability of reinforcement. Each action was reinforced randomly and independently of the animal’s action history within that trial or session. When an action was reinforced, the cue light was immediately extinguished (cue light offset) and nose pokes had no additional programmed consequences. A 0.500s delay between the final action and outcome delivery was instituted to temporally separate these events, as done in previous work (Schultz et al., 1993). Following this delay, the outcome was delivered to the animal and the food trough was illuminated. Outcomes were delivered into the food trough from a standard pellet magazine, via the operation of a smaller stepper motor and dispenser. The food trough remained illuminated and the task did not progress until the animal retrieved the outcome. Once the animal retrieved the outcome, a variable length intertrial interval (ITI) of 10-12s was initiated. In each session, 180 trials were administered.

In the first instrumental conditioning session, actions were reinforced with a probability of 1 (each action was reinforced) equivalent to a fixed ratio 1 (FR01) reinforcement schedule. In the second session, the probability that an action was reinforced was decreased across three blocks of trials. In the first block of 60 trials, actions were reinforced with a probability of 1 (FR01). In the second block of 60 trials, each action had a 1 in 3 chance of being reinforced (random ration 3, RR03). In the third block of 60 trials, the probability was further decreased to 0.2 (random ratio 5, RR05). In sessions 3 and 4, actions were reinforced with a 0.2 probability for all trials (RR05). In sessions 5-7, actions were reinforced with a probability of 0.1 for all trials (random ratio 10, RR10). In all trials but the FR01 trials, animals were required to execute an unpredictable, varying, and randomly determined number of actions per trial. Random reinforcement was utilized to limit the ability of the animal to correctly anticipate reward delivery. Actions differed from each other mainly in terms of their location within the action series in each trial (the action number within a trial, e.g. 1st action, 2nd action, 3rd action, etc.). In each trial, each animal’s action rate was calculated as the number of actions divided by the duration of the response period. This served as a measure of behavioral conditioning and performance. Changes in behavior across sessions were assessed with repeated measure analysis of variance (ANOVA), and repeated measures contrasts were applied as appropriate (Kass et al., 2014). The time interval between each action (inter-action interval) was measured for bins of low, medium, and high numbered actions (actions 1-7, 8-14, and 15-21). Statistical differences between binned inter-action intervals were assessed with repeated measures ANOVA. This measured the animal’s behavioral sensitivity to increasing action numbers throughout a trial. To examine the effects of the number of actions performed in a trial on fatigue, motivation, or attention, the number of actions performed in each trial was correlated with the latency to retrieve the reward, or initiate the next trial.

### Histology

Following the completion of experiments, animals were perfused with saline and brains were extracted. Each brain was stored in a mixture of sucrose and formalin. The brains were then frozen and sliced in 60 μ m coronal sections on a cryostat, before being stained with cresyl-violet. The location of each implant was histologically verified under light microscope according to Swanson’s brain atlas (Swanson, 2004). Animals were excluded if electrode location could not be confirmed in VTA.

### Electrophysiology

During experiments, animals were attached to a flexible headstage cable and motorized commutator that allowed the animal to move freely about the operant chamber, with minimal disruption of behavior (Plexon, Dallas, TX). Neural data were recorded via the PlexControl software package, operating a 64-channel OmniPlex recording system (Plexon, Dallas, TX). Neural data were buffered by a unity gain headstage and then a preamplifier. The digitized broadband signal was then band-pass filtered (100 Hz – 7 KHz). High-pass filtering can affect spike waveform shapes and neuronal identification, but with freely moving animals it is necessary to apply these filters to remove artifacts from the neuronal signal (Ungless and Grace, 2012). The filter pass bands that were utilized in the current manuscript are consistent with those that have previously been used to record from dopamine containing brain regions (Schultz et al., 1993; Fiorillo et al., 2003; Tobler et al., 2005). Data were digitized at 40 KHz and continuously recorded to hard disk. Voltage thresholds were applied to the digitized spike data offline (Offline Sorter, Plexon, Dallas, TX). Single units were sorted using standard techniques, and were utilized only if they had a signal to noise ratio in excess of 2/1, and were clearly separated from noise clusters and other single unit clusters. Examples of recordings and sorting are show in supplemental material (Fig. S1).

A VTA neuron was classified as dopaminergic if it had broad action potentials, greater than 1.4 ms in duration, and a mean baseline firing rate less than 10 Hz. These criteria are similar to those used in previous studies (Hyland et al., 2002; Fiorillo et al., 2003; Anstrom and Woodward, 2005; Pan et al., 2005; Tobler et al., 2005; Anstrom et al., 2007; Totah et al., 2013). All remaining neurons were classified as non-dopaminergic. All analyses were conducted on the entire population of neurons that were recorded to provide a classification-free examination of the data. Our approach to VTA cell classification is consistent with classical techniques. It should be pointed out, however, that indirect identification (using electrophysiological features as opposed genotypic features) can lead to classification errors (Margolis et al., 2006; Lammel et al., 2008; Cohen et al., 2012). In general, qualitatively similar responses were observed in both groups of neurons. Units recorded in different sessions were considered separate units, as methods to estimate neuronal identity between sessions are not widely used with VTA recordings. In total, the following numbers of neurons were recorded in each of 7 sessions: dopamine (reward responsive): 11(7), 19(10), 21(12), 29(19), 27(13), 21(9), 27(16); nondopamine (reward responsive): 31(17), 35(14), 39(27), 34(13), 25(12), 29(11), 27(9).

### Neuronal Data Analysis

Each single unit’s spike times were binned into spike counts (0.025s bins) within a trial. Binned spike counts were aligned to all events (e.g. cue light onset, actions, time period between cue light offset and outcome delivery, and outcome delivery). A stable, four s portion of the ITI (5s to 1s prior to cue light onset) served as the neuronal activity baseline. Single unit firing rates were Z-score normalized relative to baseline and zero-centered before activity was averaged together. Each unit’s normalized activity was examined in 0.250s windows around events (cue onset: +0.050 – 0.300s, relative to cue onset; -0.125 – +0.125s, relative to the time of action execution; time period between cue offset and outcome delivery: +0.150 – 0.400s, relative to execution of the last action; outcome delivery: +0.050 – 0.300s, relative to delivery). To assess between-session changes in population-level evoked activity, windowed activity was compared with a between groups two way ANOVA, with session number and neuron type (dopamine or non-dopamine) as grouping variables. In all cases, protected Fisher’s least significant difference tests were applied as appropriate.

A unit was classified as being activated or suppressed by an event if it met two criteria: 1) a significant paired samples t-test comparing raw (non-normalized) baseline firing rates with raw evoked firing rates, and 2) three or more consecutive bins of mean activity within the event-window, that were in excess of a 95% confidence interval around the baseline mean. With respect to a given task event, the proportions of units classified as being activated or suppressed were calculated. Differences in the proportions of activated units between sessions were compared with a Chi-squared test of independence. As expected, insufficient numbers of suppressed neurons (in some sessions zero) were obtained to permit reliable statistical analyses of this class of responses. Suppressed neurons are plotted for clarity, but not analyzed in detail.

The terminology “action-evoked” neuronal responses refers to activity around the time of action execution, without assuming that the action is solely responsible for evoking this neuronal response. Each unit’s activity was examined as a function of action number (a unit’s mean response to each n^th^ numbered action within a trial, across all trials). These analyses were restricted to the RR10 sessions (sessions 5-7). RR10 sessions required larger numbers of actions per trial, on average, and would ensure that there were a sufficient number of higher numbered actions (e.g. actions 18, 19, 20, 21, etc.) for analysis. All action number analyses utilized actions 1 through 21. While even higher numbered actions occurred in some trials, these actions occurred less frequently and were excluded from action number analyses, as there was insufficient sample size for reliable statistical analysis. To remove any effects of impending reward delivery on the action evoked neuronal responses, only unrewarded actions were used in this analysis. Preliminary analyses suggested that including rewarded actions had little effect on the results. Action evoked neuronal responses were binned into 3 bins of 7 consecutive actions representing low, medium, and high numbered actions (actions 1-7, 8-14, and 15-21, respectively).

We measured correlations in spike count between pairs of dopamine neurons (n = 172), pairs of non-dopaminergic neurons (n = 276), and mixed pairs (a dopamine and a non-dopamine neuron, n = 387) in RR10 sessions. Correlations were only calculated for pairs of simultaneously recorded neurons. Pairwise neuronal correlations were assessed in the same .250 sec windows used to quantify firing rates, using the standard Pearson’s correlation,

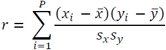

where the correlation coefficient, *r*, is the summed product of the *i*^th^ point’s residual from the sample mean, 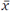 or 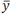, normalized by the sample standard deviation, *s*. Noise correlations represent the degree to which the joint activity of simultaneously recorded pairs of neurons fluctuates from trial to trial. Noise correlations were calculated on the trial-by-trial spike counts observed in each window. For actions, these evoked responses also examined according to action number within a trial, or binned action (actions 1-7, 8-14, or 15-21). Signal correlations refer tothe correlation between pairs of tuning curves in simultaneously recoded neurons. We corrected for chance levels of correlation, which were estimated as the mean correlation of 1000 pairs of randomly shuffled data. Fisher’s Z transformation

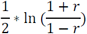

was used to transform correlation coefficients for statistical comparisons. The geometric mean spike count was calculated to measure the average joint activity of each pair of neurons.

Individual neurons preferred different subsets of action numbers. To determine the extent to which elapsed time since trial start or action number could account for action evoked neuronal activity, we performed a Poisson regression of spike count onto these variables. The percentages of neurons with significant coefficients are reported. To visualize the various tunings of VTA neurons to action number, each neuron’s mean activity as a function of action number is displayed. Because evoked firing rates could span a large range of values, each tuning curve scaled so that the maximum evoked activity was equal to 1 and the minimum evoked activity was equal to 0. Scaled tuning curves were used only for visualization. To compute the number of maxima, each tuning curve was fit with a cubic smoothing spline (smoothing parameter 0.001) and maxima were detected as points in the spline with derivatives equal to 0.

### Population Average Decoder

A decoder classified binned action number according to the trial-averaged spike count, also averaged across neurons, on an action-by-action basis. This is called the ‘population average decoder’ Binned action number, *A*, is defined as 3 bins of 7 consecutive actions (1-7, 8-14, or 15-21). The classifier maximizes

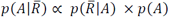

Here 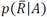 represents the probability of the population averaged response occurring, given a particular action bin. The probability of an action belonging to an action bin is denoted by *p*(A), and is uniform across action bins in the current work. That is, the prior probability for each bin, *p*(*A*), is 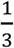. For these analyses, 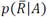 is calculated from the sample distributions of mean action evoked population responses as follows. First, the population average of all neuro’ trial averaged responses is calculated for all actions in all bins. Next, one action is selected as a test observation. Then,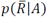 is assumed to be the best fit Gaussian of unknown mean and variance,fit from all trials in the bin *A* (excluding the test observation, if it lies in *A*). The result is 3 smooth distributions of mean activity, corresponding to the 3 bins of action numbers. The average activity of the test observations is calculated, and classified as the most likely action bin to have evoked this response, based on the relationship between population averaged activity andaction bins in the training dataset (the actions not tested). The classifier selects the action binmost likely to have evoked the population response via

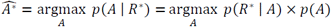

Here, *R*^∗^ is the population average of the held out action number 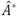 is the resulting estimatedaction bin. Testing is repeated for each action number.

### Ensemble Decoder

Assuming 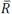 is not a sufficient characterization of 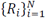, which we will call 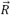,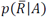, does not thoroughly represent the information in 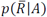. Therefore, we instead assume relevant information is captured by a higher-dimensional representation: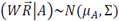.The projection, *W*, estimated by principal components analysis, is necessary to ensure the covariance, Σ, can be estimated by the usual pooled sample covariance, 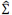 This maximizes the dimensionality of the space into which we project 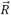, for given sample size. We then maximize 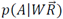. This maintains the varying firing rates in each unit, elicited by actions within different bins. Note that the population average decoder is the special case of this approach wherein *W* is a row vector of ones. The projection and classifier can be learned with each neuron’s set of evoked trial-averaged spike counts as a feature. Each action’s set of observed trial-averaged spike counts, the new test observations, 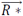, are then projected via the weights estimated from the training data and classified sequentially (that is, we estimate the set of 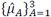 on all the remaining action numbers, which is equivalent to training, and maximize the analogous posterior).

### Statistical Testing of the Decoders

To test whether the two decoders produce equal or different classification accuracy, we fit a binomial generalized linear model and controlled for action number, session, and unit type (dopaminergic versus non-dopaminergic). For each trial of action bin *A*, tested on decoder *d*, the log odds of the probability of correct classification was assumed to be linear in its covariates as given by the logistic regression function

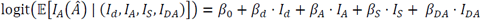

Where 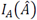 is an indicator signifying correct or incorrect classification, and *I*_*d*_, *I*_*A*_, *I*_*S*_, *I*_*DA*_ are indicators (or a set of indicators) for the decoder, action bin, session, and dopaminergic unit, respectively. Likelihood ratio tests were used to test the effect of each group of factors, corresponding to each feature. Additionally, statistical significance of decoder performance (versus chance levels of correct classification) at each action number was determined via permutation test. Approximations to the exact p-values can be determined by assigning classes to each action according to a uniform prior distribution, and calculating the proportion of sets of resulting classifications that perform better than the classification rates observed using the previously discussed methods.

## RESULTS

### Animals learn to execute serial actions for reward

VTA activity was recorded while rats learned to execute multiple actions for random reinforcement with one sugar pellet (Fig. 1A, B). In the first session, each action was rewarded (FR01). In session 2, the reward probability was decreased from 1 to 0.2 across 3 blocks of trials. In sessions 3 and 4, actions were reinforced at a probability of 0.2 (RR05). In sessions 5 – 7, actions were reinforced at a probability of 0.1 (RR10). In all randomly reinforced trials, unpredictable and varying numbers of actions were required, but each action was equally likely to be reinforced.

**Fig. 1.**
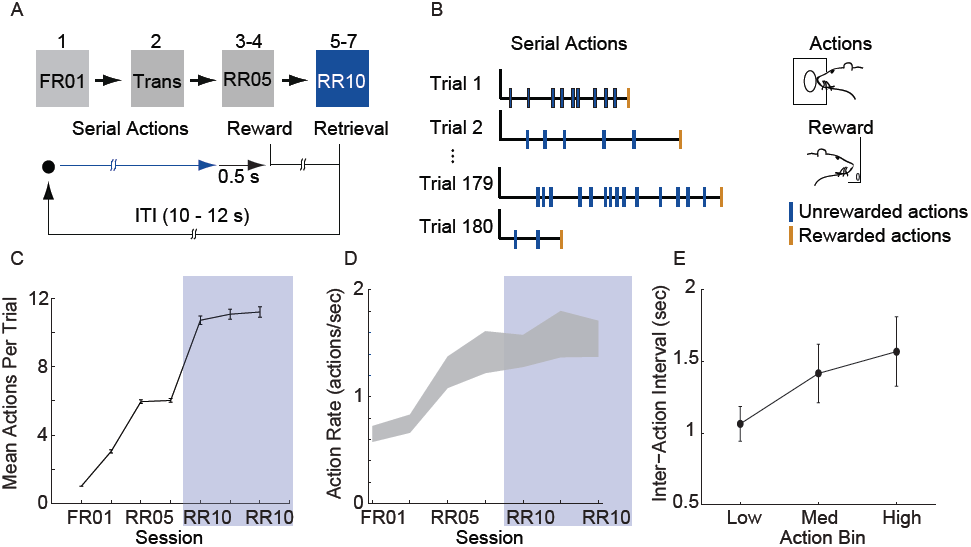
Animals learn to perform serial actions. (A) Actions (nose pokes into a lit port) were reinforced probabilistically with sucrose pellets (rewards). In session 1, each action was reinforced (fixed ratio 1, FR01). In session 2, reinforcement probability decreased to 0.2 in 3 trial blocks (transition, TRANS). In sessions 3-4, the probability of reinforcement was 0.2 (random ratio 5, RR05). In sessions 5-7, the probability of reinforcement was 0.1 (random ratio 10, RR10). At trial start, a cue light was illuminated until the outcome was earned (blue arrow). When an action was reinforced, the cue light was extinguished immediately and the outcome was delivered 0.5 sec later. (B) Random reinforcement led to different numbers of serial actions performed per trial. (C) Mean ± SEM number of actions required per trial. Shaded area depicts RR10 sessions. Serial action data were drawn from these sessions because these sessions required the greatest average number of actions per trial, and thus, had the greatest statistical power. (D) Mean ± SEM action rate in each session. (E) Mean ± SEM inter-action intervals in low, medium and high bins (actions 1-7, 8-14, and 15-21) during RR10 sessions.

Action response rates increased as reinforcement probability decreased (Fig.1C, D), and the response rate in the final RR10 sessions was significantly higher than all other sessions (Fig. 1D; F_(6,24)_ = 4.726, p = 0.003). To investigate a potential relationship between behavioral performance and action number, we divided the RR10 data into three bins: actions 1-7 (low), actions 8-14 (medium), and actions 15-21 (high). We analyzed serial actions in trials reinforced with a probability of 0.1 (RR10 sessions, sessions 5-7), because this reinforcement schedule produced the greatest average number of actions performed per trial (Fig. 1C). Within a trial, the inter-action interval significantly increased with increasing action number bins (Fig. 1E; F_(2,44)_ =11.020, p < 0.001). This indicated that behavior was sensitive to action number. The latency to retrieve the reward (r_(3468)_ = -0.017, p = 0.309) or initiate responding in the next trial, however, was not correlated with binned action number (r_(3445)_ = 0.015, p = 0.389).These data suggest that the number of actions performed in a trial did not affect behavioral responses to reward or cues.

### Neurophysiological recordings

A total of 375 units (in 7 sessions and from 10 rats) were recorded from electrodes histologically localized to VTA (Fig. 2). All neurons were included in these analyses because we were interested in understanding the full diversity of activity patterns in the VTA. Neurons were classified (Fig. 3) as putative ‘dopamine’ (n = 155) or ‘non-dopamine’ neurons (n=220) based on electrophysiological criteria. This classification approach permits comparison with previous work, despite potential inaccuracies (Margolis et al., 2006) and with the caveat that some dopamine neurons co-release other neurotransmitters (Tritsch et al., 2012). We did not observe strong clustering in the electrophysiological profiles of these neurons. It is important to underscore that neurons were collected with no attempt to select for particular electrophysiological characteristics. When VTA neurons are recorded in an unbiased manner, as done here, weak clustering may be expected (Kiyatkin and Rebec, 1998). In addition to the above classification, analyses were performed on reward-responsive dopamine neurons, as this subgroup may be a more conservative estimate of dopamine neuron identity (Lak et al., 2014; Eshel et al., 2016).

**Fig. 2.**
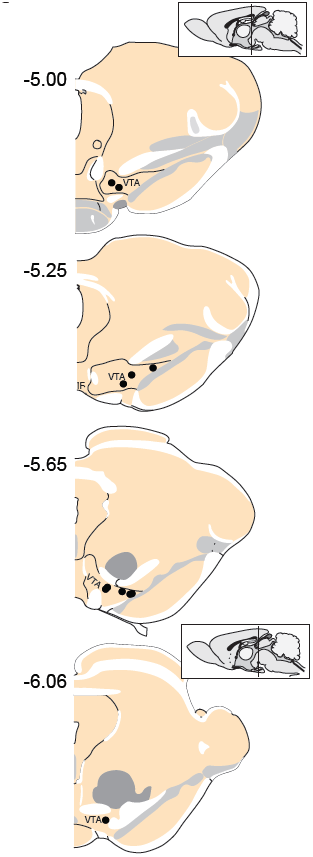
Locations of recording electrodes within VTA. Each dot represents the location of a recording array. Insets show midsagittal diagram of rodent brain with vertical lines representing the approximate location of the most anterior and posterior coronal sections shown.

**Fig. 3.**
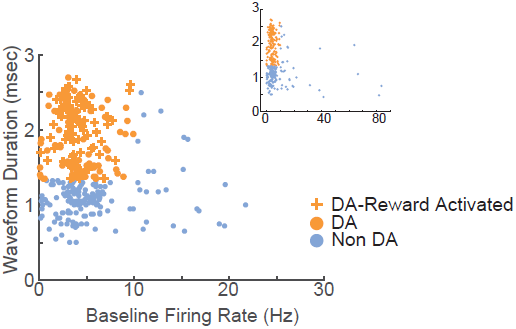
Electrophysiological characterization of recorded VTA neurons. Each point represents a single neuron; data from all sessions are depicted. Neurons with wide spike widths (>1.4 ms) and low firing rates (< 10 Hz) were considered putative dopamine neurons. Crosses depict reward-responsive dopamine neurons. The remaining neurons were considered putative non-dopamine neurons. The main figure depicts the data with the horizontal axis limited to 30, for better visualization of the majority of the data. Inset shows the same data, with the horizontal axis extended to encompass the full range of baseline firing rates.

### VTA neurons are diversely tuned to serial actions

In order to understand how VTA neurons encode information during serial actions, we examined how activity was modulated by actions in RR10 sessions. Action evoked responses initiated -0.173±0.013 sec prior to actions and had an average duration of 0.313±0.012 sec (Fig. 4A). Thus, the majority of neurons had response durations several hundred ms faster than the majority of inter-action intervals, and therefore, action-evoked responding was well separated between successive actions in a trial. When RR10 actions were examined according to their number within a trial (e.g. 1^st^, 2^nd^, 3^rd^ action in a trial), we observed that individual neurons fired selectively for subsets of actions (Fig. 4B, C). We observed preferential responding to low, medium, or high numbered actions within the population of simultaneously recorded neurons (Fig. 4D-F). To determine how VTA neurons responded to serial actions, we calculated each neuron’s tuning curve as a function of action number within a trial (Fig. 4G). Action tuning curves mostly had a single peak (local maxima, χ^2^_(1)_= 9.921, p = 0.002; neuron type, χ^2^_(1)_=1.267, p = 0.260), and different neurons preferred different subsets of actions (Fig. 4G,H). Tuning curve depth (maximum – minimum) did not differ significantly between putative dopamine and non-dopamine neurons (Fig. 4I; t_(154)_ = 3.224, p = 0.084). Averaging activity across neurons, a traditional approach to analyzing VTA neuronal data, concealed this heterogeneity in both groups of neurons (Fig. 4J).

**Fig. 4.**
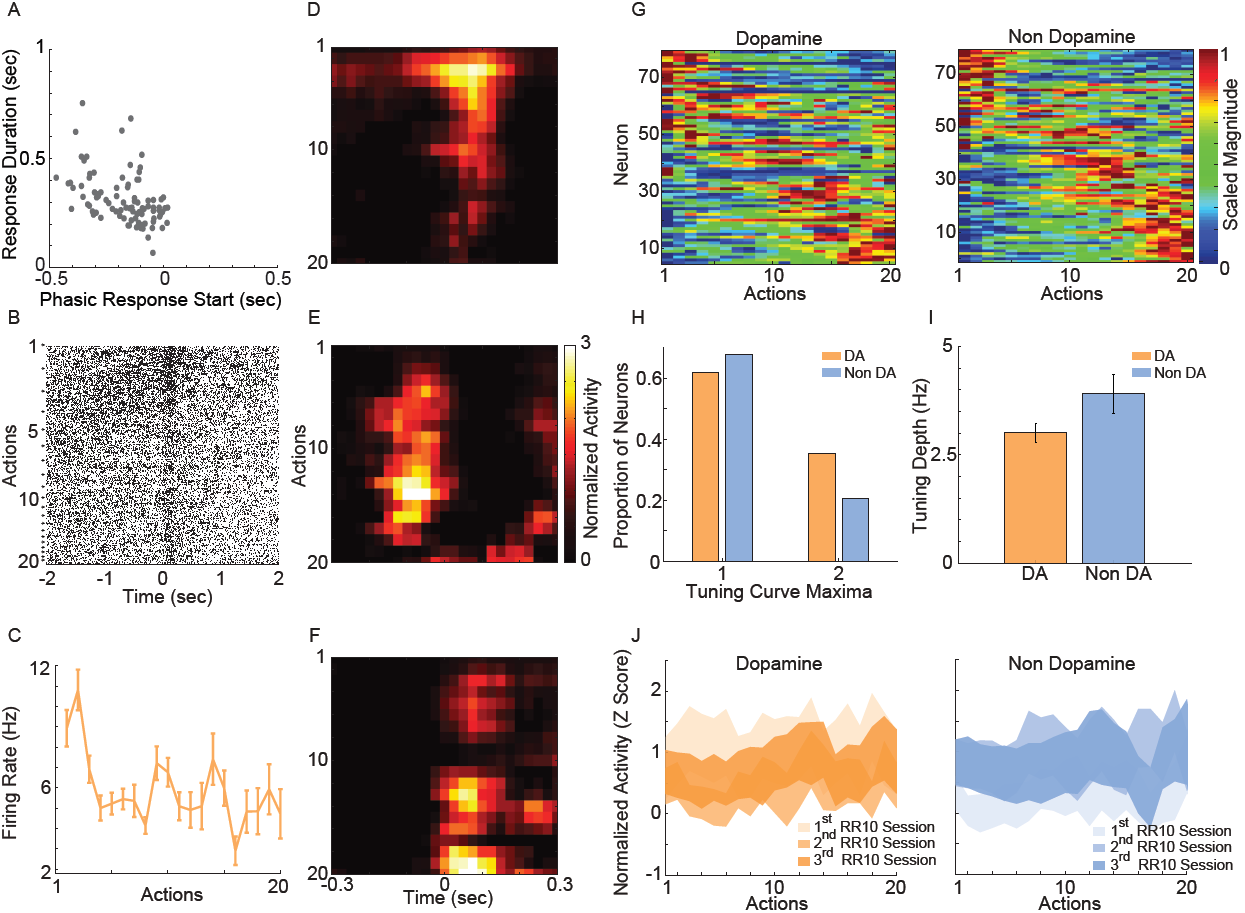
RR10 action evoked VTA responses. (A) Scatter plot depicts response start and duration of action evoked responses in RR10 sessions. Each point represents the data from a single unit. (B) Data from a typical dopaminergic neuron that preferred low action numbers. The raster shows spikes aligned to the time of action execution (time = 0). Each row of the raster represents one action evoked response and rows are arranged by action number. Each arrow on the right represents an action number. For each action number, the earlier occurrences of an Nth numbered action are arranged toward the top. Thus, the first row of the raster represents the first occurrence of an action number 1 (i.e. trial 1), and the second row represents the second occurrence (i.e. trial 2), etc. (C) Depicts the action-evoked responses of the neuron depicted in (B) conditioned on action numbers 1-20. Note that the neuron most strongly prefers actions 1 and 2. (D-F) Neuronal responses aligned to actions for 3 simultaneously recorded neurons. (D) A non-dopaminergic neuron that fires preferentially during lower numbered actions within a trial. Data depicted as the average response evoked by actions 1-20. Data are aligned to the time of action execution (0.025 sec bins). Only unrewarded actions are depicted. (E-F) A pair of dopaminergic neurons that fire preferentially during upper-middle, and higher numbered actions, respectively. Note that both dopaminergic and non-dopaminergic VTA neurons fire preferentially around distinct subsets of actions.(G) Action number tuning curves during RR10 (0.25 sec window, centered on action). Each dopamine (left) and non-dopamine (right) neuron’s average, action evoked, normalized, firing rate during execution of actions is plotted by color. Tuning curves are scaled to each curve’s peak (hot colors = tuning curve peak). Neurons are sorted by the location of the peak of the tuning function. Note the diversity of tuning curves to serial actions (H) Proportion neurons with 1 or 2 tuning curve maxima. (I) Mean ± SEM tuning curve depth. (J) Mean ± SEM neuronal responses as a function of action number in RR10 sessions for dopamine and non-dopamine neurons.

The diversity of the tuning curves of VTA neurons suggested that network properties may be critical to information processing during serial actions. To gain insight into VTA network-based information processing, we calculated the trial-by-trial correlations in spike counts (noise correlations), and correlations between tuning curves (signal correlations) for all simultaneously recorded pairs of neurons. Noise correlations reflect functional connectivity between neurons while signal correlations reflect similarity in tuning curves (Cohen and Kohn, 2011). Correlations were examined between all possible pairings of VTA neurons (dopamine – dopamine, non-dopamine – non-dopamine, and dopamine – non-dopamine).

We found that noise correlations during actions were significantly lower in pairs of nondopamine neurons than pairs containing a dopamine neuron (Fig. 5A; F_(2,858)_ = 4.052, p = .018). The magnitude of these correlations was decreased during low numbered actions compared to medium or high numbered actions (F_(2,1716)_ = 14.625, p < .001). It is unlikely that these differences were due to differences in spike count because the pairwise geometric mean spike count was not associated with noise correlation magnitude in any bin (low: r = 0.008, p = .805; med: r = -0.015, p = .650; high: r = -0.048, p = .158). In contrast, signal correlation strength did not differ between different pairings of VTA neurons (Fig. 5B; F_(2,858)_ = 0.289, p = .749). There was a strong association between signal correlations and noise correlations (Fig. 5B, Table 1), suggesting that there is a high degree of functional connectivity between pools of similarly tuned neurons. This result raises the possibility that different patterns of functional connectivity contribute to the diversity of tunings to actions.

**Table 1.** Table depicts the correlation between action number signal correlations and action evoked noise correlations. Data are taken from RR10 sessions. Data depicted separately for each pair type (DA-DA, dopamine pair; DA-NDA, dopamine and nondopamine pair; NDA-NDA, non-dopamine pair).

|  | Rho | p value |
| --- | --- | --- |
| DA – DA Pair | 0.571 | < 0.001 |
| DA – NDA Pair | 0.569 | < 0.001 |
| NDA – NDA Pair | 0.528 | < 0.001 |

**Fig. 5.**
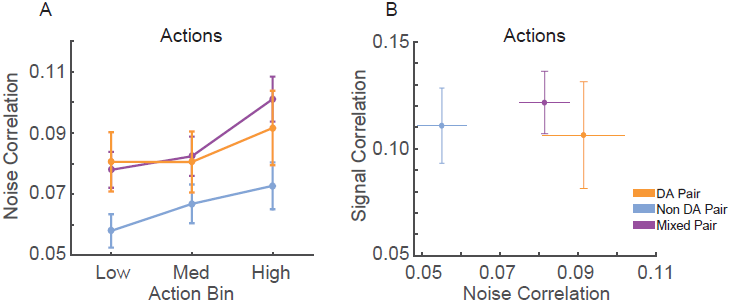
Correlation structure of RR10 action-evoked responses. (A) Mean ± SEM action evoked noise correlations between simultaneously recorded pairs of VTA neurons in RR10 sessions during actions. Noise correlations measure trial-by-trial fluctuations in pairwise activity. Data depicted as mean ± SEM noise correlation in each bin of actions. Data are plotted separately for all pairings of VTA neurons. (B) Mean ± SEM signal correlation (correlation between tuning curves) and noise correlations during RR10 actions. For simplicity, noise correlations are collapsed across action number bin.

### Actions in a trial are accurately discriminated from VTA neuronal ensemble activity

The combination of tuning curve heterogeneity, lack of population average signal for serial actions, and the association between action-evoked signal and noise correlations suggested that information about ongoing sequences of actions may be encoded by VTA ensembles. To quantify the accuracy of this encoding mechanism, we differentiated low, medium, and high action number bins from either population-averaged activity (Fig. 6A) or the collective activity of VTA ensembles (Fig. 6B). The same cross-validated linear discriminant analysis decoding algorithms were used for both the ensemble and population-average data (see Methods). Ensemble activity was discriminated more accurately than shuffled control data (Fig. 6C; 85% accuracy versus chance levels of correct decoding [33% accuracy], permutation test, p < 0.001). In contrast, population averaged activity was decoded at chance levels (Fig. 6C; permutation test, p = 0.240). Binned action number was discriminated from ensemble activity significantly more accurately than population-averaged activity (Fig. 6C; χ^2^_(1)_= (21.435, p < 0.001). Discrimination was stable across sessions (χ^2^_(2)_= 1.097, p = 0.578), and between low, medium, and high action number bins (χ^2^_(2)_= 3.614, p = 0.164). There was no difference in accuracy between dopamine and non-dopamine neurons (χ^2^_(1)_= 1.133, p = 0.287). When reward responsiveness was also included in the model, dopamine and non-dopamine neurons were decoded at similar accuracy (χ^2^_(1)_= 1.300, p = 0.254). These data suggest that real-time information about ongoing behavior is encoded by the collective activity of VTA neurons, but not the averaged activity of VTA dopamine or non-dopamine neurons. Thus, VTA ensemble activity accurately predicted the current action bin, and could be used to organize, pace, and sustain effortful behaviors.

**Fig. 6.**
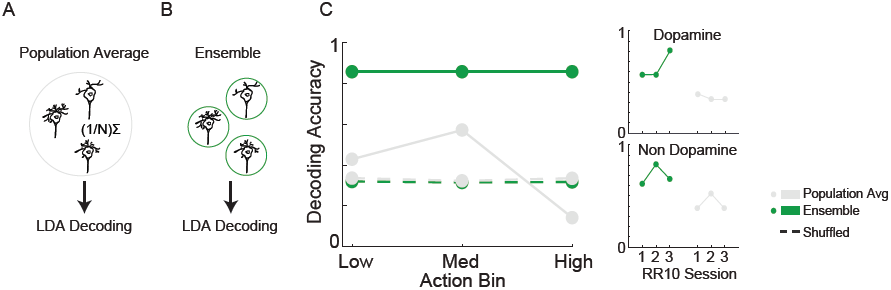
Decoding binned action number from VTA neuronal activity. Linear discriminant analysis (LDA) utilized action-evoked activity averaged across (A) all neurons (population-average), or (B) the collective activity of each neuron (ensemble). (C) The proportion of correctly classified actions for each decoder. Dashed lines represent decoding of shuffled control data. Inset to the right depicts performance of ensemble and population-average decoders across consecutive sessions in dopamine and non-dopamine neurons. Note that ensemble activity was decoded significantly more accurately than population averaged activity and shuffled control. Inset data are collapsed across high, medium, and low action numbers and depicted separately for each session.

To determine if serial actions or the passage of time preferentially modulated neuronal activity, we examined the relationship between action evoked spike counts and action number, or the corresponding time within a trial. Action number predicted spike counts of a significantly larger proportion of neurons than time elapsed in a trial (χ^2^_(1)_= 20.643, p < 0.001; 29% versus 7% of neurons, respectively). Actions did not preferentially modulate the activity of either dopamine or non-dopamine neurons (χ^2^_(1)_= 0.179, p = 0.673). These data suggest action-evoked activity reflected ongoing behavior and do not support a role for encoding the passage of time in this task.

Neuronal responses may have represented reward anticipation (Bromberg-Martin et al., 2010; Totah et al., 2013) in a manner which changed throughout a series of actions. If this were the case, the number of actions performed in a trial would be correlated with neuronal activity in a consistent direction. Thus, we examined the correlation between neuronal responses and the number of actions in each trial. Only a small proportion of dopamine and non-dopamine neurons had significant correlations between the response evoked by the first action in a trial and the number of actions in the previous trial (Table 2). Similar proportions of these neurons were positively or negatively correlated, indicating that this relationship was inconsistent between neurons (Table 2). Likewise, very few neurons had responses to the last action in each trial that were significantly correlated with the number of actions in that trial; there was no significant difference in the proportion of dopamine and non-dopamine neurons with positive or negative correlations (Table 2).

**Table 2.**
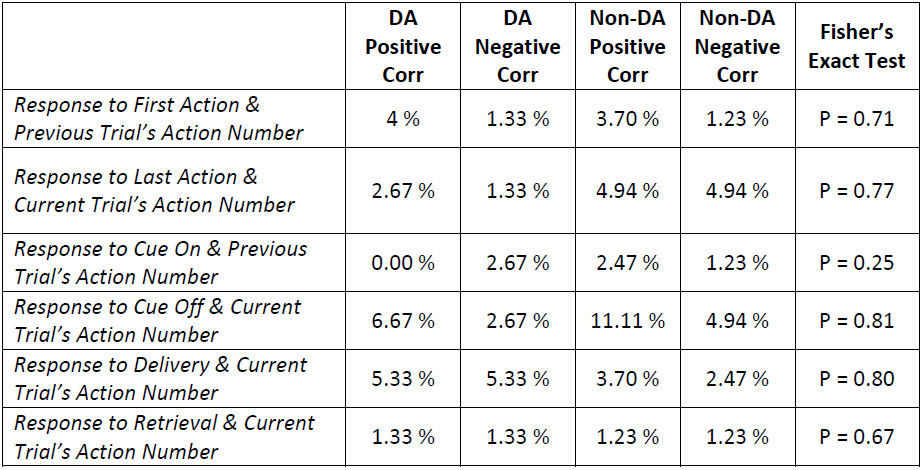
Table depicts the percentage of neurons with significantly correlated responses between task events and the number of actions in each trial. Data are taken from RR10 sessions. Pearson’s correlations were calculated on spike counts taken from 0.25s windows around events. The far right column depicts the p value corresponding to a Fisher’s exact test of these proportions.

|  | **DA**<br>**Positive**<br>**Corr** | **DA**<br>**Negative**<br>**Corr** | **Non-DA**<br>**Positive**<br>**Corr** | **Non-DA**<br>**Negative**<br>**Corr** | **Fisher's**<br>**Exact Test** |
| --- | --- | --- | --- | --- | --- |
| *Response to First Action &*<br>*Previous Trial's Action Number* | 4 % | 1.33 % | 3.70 % | 1.23 % | P = 0.71 |
| *Response to Last Action &*<br>*Current Trial's Action Number* | 2.67 % | 1.33 % | 4.94 % | 4.94 % | P = 0.77 |
| *Response to Cue On & Previous*<br>*Trial's Action Number* | 0.00 % | 2.67 % | 2.47 % | 1.23 % | P = 0.25 |
| *Response to Cue Off & Current*<br>*Trial's Action Number* | 6.67 % | 2.67 % | 11.11 % | 4.94 % | P = 0.81 |
| *Response to Delivery & Current*<br>*Trial's Action Number* | 5.33 % | 5.33 % | 3.70 % | 2.47 % | P = 0.80 |
| *Response to Retrieval & Current*<br>*Trial's Action Number* | 1.33 % | 1.33 % | 1.23 % | 1.23 % | P = 0.67 |

We further investigated if reward anticipation modulated neuronal responses by examining the relationship between responses evoked by the cue light turning off, reward delivery, or reward retrieval, and the number of actions performed in each trial. We found no significant differences in the proportion of dopamine or non-dopamine neurons with positive or negative correlations between responses evoked by cue onset and the number of actions in the previous trial or between cue light offset, reward delivery, or reward retrieval responses and the number of actions in the current trial (Table 2). Taken together, these results indicate that performance of serial actions did not influence encoding of reward anticipation, supporting the notion that VTA neurons encoded information about ongoing behaviors as opposed to information about time or reward anticipation in the current task.

### Reward delivery evoked responses

Reward delivery evoked greater activity in dopamine neurons than in non-dopamine neurons (Fig. 7A, B; F_(1,361)_ = 9.159, p = .003). This pattern did not change across sessions (Fig. 7C; F_(6,361)_= 1.352, p = .233). The proportion of dopamine neurons responding to reward delivery did not vary significantly across sessions (Fig. 7D; all neurons, X^2^_(6)_= 9.289, p = .158; non dopamine, X^2^_(6)_= 13.234, p = .039).

**Fig. 7.**
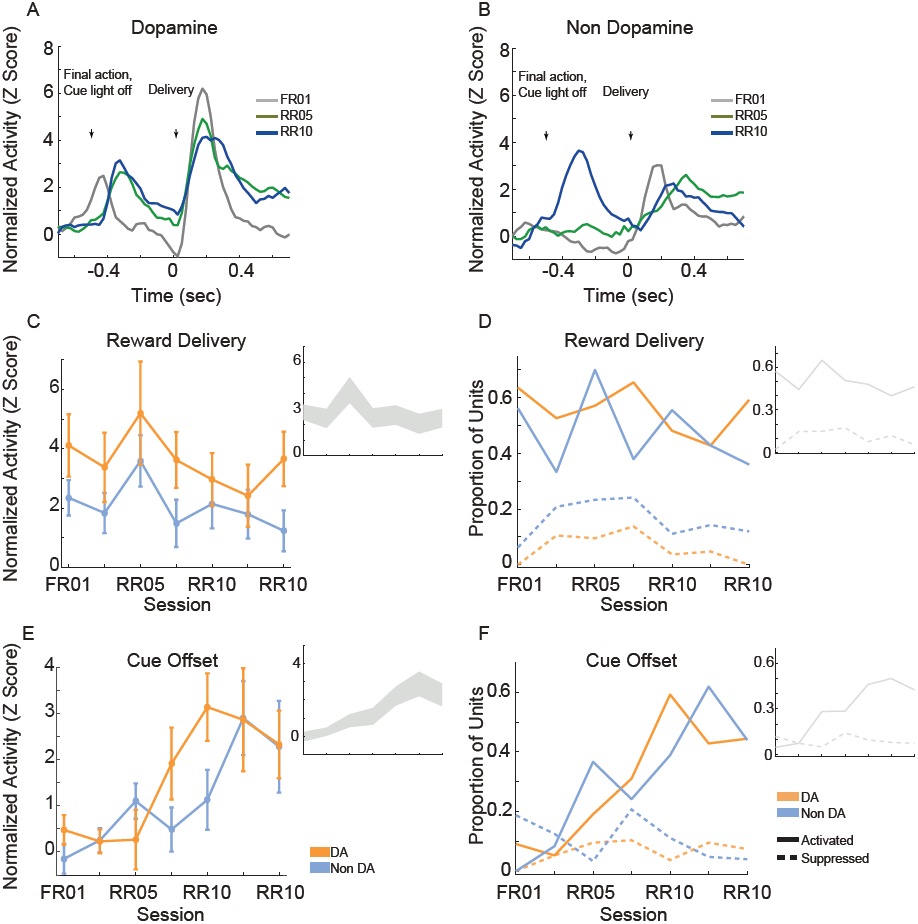
VTA responses to rewards and reward predictive cues. Normalized mean (A) dopamine and (B) non-dopamine responses aligned to outcome delivery (time = 0). The final action in each trial (left arrow) occurred 0.5s prior to outcome delivery (right arrow). Cue light offset was simultaneous with execution of the final action, which signaled trial completion and impending outcome delivery. Data are depicted for FR01, final RR05 session, and the final RR10 session. (C) Mean ± SEM responses evoked by reward delivery (+0.05 - +0.3s, relative to reward delivery) in both groups of neurons and all sessions. Dopamine responses were greater than non-dopamine neuronal responses. Inset depicts data from all VTA neurons plotted without respect to neuronal classification. (D) The proportion of neurons classified as either significantly activated (solid lines) or suppressed (dashed lines) by outcome delivery. Data are depicted across all sessions for putative dopamine and non-dopamine neurons. Inset depicts data from all VTA neurons plotted without respect to neuronal classification. (E) Mean ± SEM responses following cue offset, during the pre-reward delivery period (+0.150 - +0.400s, relative to cue offset). Data are depicted similarly to (C). In each group, the evoked response increased across sessions. (F) The proportion of neurons activated (solid lines) or suppressed (dashed lines) by cue offset signaling future reward delivery. Data depicted similarly to (D).

After the final action in a trial, the cue light was immediately extinguished and reward was delivered 0.5 s later. We hypothesized that cue light offset would evoke VTA responses, similar to a reward prediction error, because this event signaled new information about unpredictable outcome delivery (Montague et al., 1996; Schultz, 1998). As training progressed, cue offset evoked great activity levels (Fig. 7E; F_(6,361)_= 4.776, p < .001), and activated a greater proportion of neurons (Fig. 7F; all neurons, X^2^_(6)_ = 45.949, p < .001; dopamine neurons, X^2^_(6)_ =22.093, p = .001; non dopamine neurons, X^2^_(6)_= 29.744, p < .001). This finding confirms that we observed responses to cues and rewards that are commonly attributed to dopamine neurons (Montague et al., 1996; Schultz, 1998), but also occur in non-dopaminergic cells in the VTA (Kim et al., 2010; Cohen et al., 2012; Kim et al., 2012).

### Cue evoked responses

When animals were required to perform a single action to earn rewards (FR01), the majority of dopamine neurons encoded cue onset (Fig. 8A), which is consistent with previous observations (Schultz, 1998; Roesch et al., 2007). Non-dopamine neurons followed a similar pattern (Fig. 8B). During random ratio sessions, the population response to the cue decreased significantly as response requirement increased (Fig. 8C; session, F_(6,361)_ = 2.667, p = .015). This pattern of responding did not differ between dopamine and non-dopamine neurons (Fig. 8C; F_(1,361)_ = 1.543, p = .215). We found the same effects when analysis was restricted to neurons that were significantly activated by reward delivery (session, F_(6,175)_ = 4.961, p < .001; neuron type, F_(1,175)_ = 3.912, p = 0.050). The data among neurons not responsive to reward delivery were inconclusive, but followed the same trend (session, F_(6,172)_ = 2.096, p = 0.056; neuron type, F_(1,172)_ = 0.824, p = 0.365). Likewise, significantly fewer neurons were modulated by cue light onset in random ratio sessions (Fig. 8D; all VTA, X^2^_(6)_= 23.844, p = .001; dopamine, X^2^_(6)_ =20.109, p = .003; non-dopamine, X^2^_(6)_= 10.636, p = .100).

**Fig. 8.**
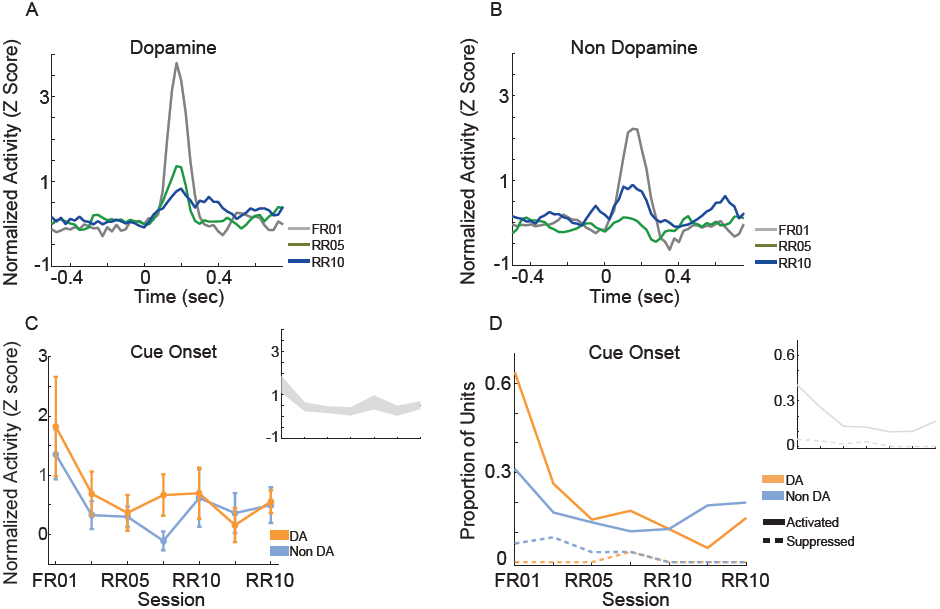
VTA responses to cue onset. Normalized mean (A) dopamine and (B) non-dopamine responses to cue onset (time = 0). Data are depicted for the FR01 session, the final RR05 session, and the final RR10 session. (C) Mean ± SEM responses evoked by cue onset (+0.05 - +0.3s) in all groups of neurons and sessions. Data are depicted separately for all putative dopamine and non-dopamine neurons. Responses were greatest in session 1 and significantly less in subsequent sessions, with no difference by neuron type. Inset depicts data from all VTA neurons plotted without respect to neuronal classification. (D) Proportion of neurons activated (solid lines) or suppressed (dashed lines) by cue onset. Data are depicted across all sessions for putative dopamine and non-dopamine neurons. The proportion of activated dopamine neurons, but not non-dopamine neurons, decreased in later sessions. Inset depicts data from all VTA neurons plotted without respect to neuronal classification.

### Cue and reward evoked responses are decoupled when performing serial actions

We assessed the correlation between each neuron’s responses to cue onset and reward delivery in order to understand the relationship between encoding of these events. During FR01 sessions, responses evoked by these events were significantly correlated in both dopamine and non-dopamine neurons (Table 3). This correlation decreased in RR05 sessions (Table 3), and was no longer significant by RR10 sessions (Table 3). A similar pattern was observed in dopamine neurons that were activated by reward delivery (Table 3). This suggests that, although cue and reward responses are positively correlated in Pavlovian conditioning tasks (Eshel et al., 2016), cue-evoked responses become decoupled from reward responses during an instrumental random reinforcement schedule. Figure 9 depicts two example neurons in the FR01 session (Fig. 9A) and the final RR10 session (Fig. 9B). Note that this example neuron has similar responses to cue onset and reward delivery in the FR01 session (Fig. 9A), while the example neuron (same animal) recorded during the RR10 session does not respond to cue onset, but is activated by reward delivery (Fig. 9B).

**Table 3.**
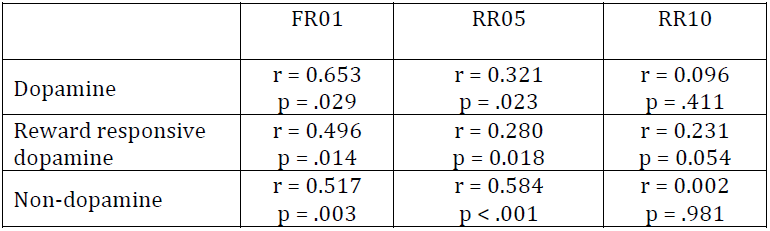
Table depicts the correlation between cue onset and reward delivery evoked responses. Data depicted separately for dopamine neurons, reward responsive dopamine neurons, and non-dopamine neurons, in FR01, RR05, and RR10 sessions. Note that the correlation between cue and reward delivery evoked response magnitudes decreases from FR01 to RR10 sessions.

**Fig. 9.**
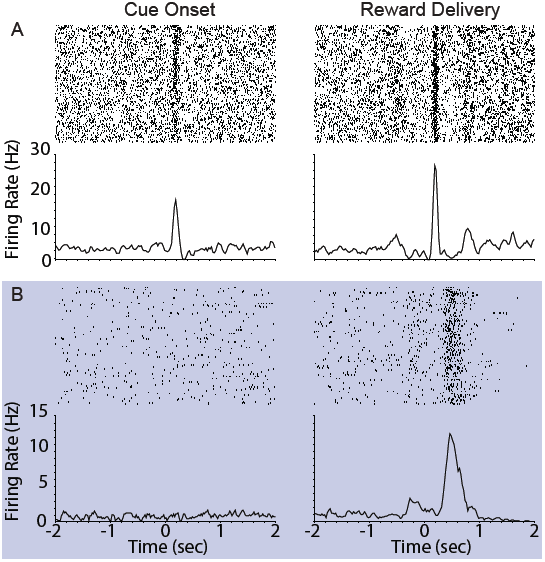
Example VTA responses to cue onset and reward delivery in first session (FR01) and final RR10 session. (A) Top left panel depicts representative example raster of cue evoked response (cue onset and time =0; trials arranged in chronological order starting at top). Each dash represents one spike. Smoothed mean response is depicted below raster. The same neuron’s response to reward delivery is depicted in the upper right panel. Note the phasic response to both cue onset and reward delivery in the FR01 session (top row). (B) Representative example neuronal responses recorded in final RR10 session (data recorded from same animal in panels (A) and (B)). Data depicted similarly to (A) with different Y axis on bottom panels. Note that this neuron responds to reward delivery, but not cue onset.

### Modulation of noise correlations by cue onset and reward delivery in serial action trials

We calculated the signal and noise correlation between pairs of VTA neurons during cues and reward delivery in RR10 sessions. We found a significant interaction between pair type (dopamine pairs, non-dopamine pairs, and mixed pairs) and event (Fig. 10; F_(4,2496)_ = 9.482, p < .001). There was no difference in baseline noise correlations between any pairing of VTA neurons (Fig. 10; F_(2,832)_ = 0.855, p = .426). Noise correlations evoked by cue onset were significantly greater in pairs containing a dopamine neuron than pairs of non-dopamine neurons (Fig. 10; F_(2,832)_ = 5.881, p = .003). There was no association between cue evoked noise correlations and pairwise cue evoked geometric mean spike counts (r = 0.008, p = .817), suggesting that mean activity level did not account for this difference. Noise correlations were strongest during reward delivery overall. Noise correlations during reward delivery did not differ significantly between pairings of neurons (Fig. 10; F_(2,832)_ = 2.794, p = .062). These data confirm previous findings that correlated activity in VTA circuits emerges when rewarding outcomes can be earned (Joshua et al., 2009; Kim et al., 2012; Eshel et al., 2016).

**Fig. 10.**
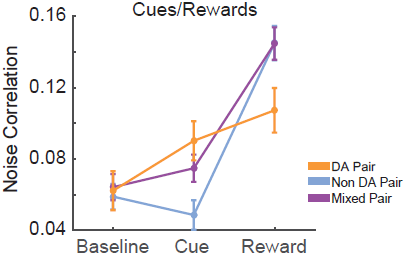
Correlation structure of cue and reward evoked responses. Mean ± SEM noise correlation between simultaneously recorded pairs of VTA neurons in RR10 sessions, during baseline, cue onset, and reward delivery. Data depicted separately for all pairings of neurons. Note that reward delivery evoked the largest magnitude noise correlations in all pairings.

## DISCUSSION

We investigated how VTA neurons encode information when animals execute an uncertain numbers of actions to earn a reward. Dopamine and non-dopamine VTA neurons were preferentially tuned to different subsets of binned action number in a manner that allowed serial actions to be accurately discriminated from the collective activity of VTA ensembles. In addition, pairs of neurons with similar action tuning curves had higher action-evoked noise correlations suggesting that VTA network properties may be critical to information processing during serial actions. Our analyses further showed that this pattern of tuning is distinct from reward prediction related activity or the passage of time. We also demonstrated that cues preceding a series of actions, unlike a single rewarded action, do not evoke neuronal responses and are decoupled from reward, suggesting that cue-related information is utilized differently when a single versus an unpredictable series of actions are required to obtain reward.

### Encoding information during an action series

Previous reports indicate that dopamine neurons encode errors in the predicted value of rewards as well as action values (Montague et al., 1996; Schultz, 1998; Morris et al., 2006; Roesch et al., 2007). Further, dopamine neurons phasically fire during the first and last action in a sequence, which may be critical to initiating and terminating behavior (Jin and Costa, 2010). Our data indicate that the collective activity of ensembles of VTA neurons also has the capacity to encode information about ongoing behavior in real-time.

The ensemble code revealed here may be used to organize and sustain actions, which is a fundamental feature of dopamine’s role in cognition (Aberman and Salamone, 1999; Ishiwari et al., 2004; Salamone et al., 2009). In particular, this signal could be critical for supporting the unique motivational and cognitive demands when large numbers of actions are required (Goldman-Rakic, 1998; Aberman and Salamone, 1999; Salamone and Correa, 2002; Seamans and Yang, 2004; Robbins and Roberts, 2007; Salamone et al., 2007; Robbins and Arnsten, 2009). VTA dopamine neurons project to several networks that may utilize this signal to organize behavior. For instance, dopamine innervation of the ventral striatum is selectively required for completing a lengthy series of instrumental actions, but not for executing small numbers of actions for reward (Aberman and Salamone, 1999; Ishiwari et al., 2004; Mingote et al., 2005). Dopamine projections to the prefrontal cortex are important for working memory and related constructs that require real-time information about ongoing behavior (Goldman-Rakic, 1998; Seamans and Yang, 2004; Robbins and Roberts, 2007; Robbins and Arnsten, 2009). VTA neurons could contribute to these functions by continually encoding information throughout execution of a lengthy series of actions, allowing striatal and prefrontal networks to track goal-directed effort expenditure and progress.

Our analyses suggest that VTA responses do not represent the passage of time in the current task. Although dopamine responses are sensitive to the timing of reward delivery and necessary for perception of time and interval timing (Lake and Meck, 2013; Bermudez and Schultz, 2014; Parker et al., 2015; Soares et al., 2016), neuronal responses were not consistently modulated by time elapsed in a trial. Instead, activity during a series of actions was most strongly modulated by action execution. This is likely because reward delivery was not contingent upon elapsed time, and VTA neurons are sensitive to the contingency between action execution and outcome delivery. Experimental designs with an explicit contingency between elapsed time and reward delivery, such as a fixed interval reward schedule, may reveal a more complex relationship between action execution and within trial timing in the responses of VTA neurons.

We also found that the number of actions required in RR10 trials did not modulate VTA responses to the rewarded action or the first action in the next trial. This suggests that recent trial history does not affect action evoked responses at the beginning or end of a series of actions. Similarly, VTA responses to stimuli associated with reward were not consistently correlated with the number of actions that an animal performed, suggesting that VTA correlates of reward anticipation did not increase or decrease according to action number. This result was expected, as each action was reinforced with equal probability.

### VTA ensemble encoding

Neurons with similar action number tuning curves had higher action-evoked noise correlations in spike count, which reflect shared connectivity between neurons (Cohen and Kohn, 2011). This suggests that different action number tunings may arise from unique inputs or differing connection strengths between inputs and VTA neurons with similar tunings may share connection properties. VTA receives inputs from an expansive set of afferent structures (Geisler and Zahm, 2005; Geisler et al., 2007), and the diversity of inputs that converge in VTA may contribute to preferential firing in different subsets of serial actions.

The association between noise correlations and action selectivity indicates that VTA network properties are critical to understanding how information about serial behavior is encoded. Networks can encode information redundantly or through diverse activity patterns and heterogeneous tunings naturally lead to the capacity to encode information as ensembles. Accordingly, serial actions were accurately discriminated from ensemble activity, but not from population-averaged activity. Though previous studies suggest VTA activity is highly redundant (Joshua et al., 2009; Schultz, 2010; Glimcher, 2011; Kim et al., 2012; Eshel et al., 2016), this work was limited to tasks requiring very few actions. The diversity of VTA tuning curves may increase to match the expanded behavioral state space of serial actions compared with the limited state space of single action trials (Eshel et al., 2016). Thus, ensemble encoding of information may occur when the complexity of a task increases, such as during ongoing serial actions.

Units classified as either putative dopamine or non-dopamine neurons had comparable mixed tunings to serial actions, suggesting that dopamine neurons can cooperate with nondopamine VTA neurons to encode information about serial actions. These ensemble signals consisting of multiple neurotransmitters could allow information to be decoded through multiple signaling mechanisms, which may diversify the spatiotemporal properties of this signal (Seamans and Yang, 2004; Kim et al., 2010; Barker et al., 2016). The activity of different types of VTA neurons must ultimately be coordinated and unified to represent information, and the present work demonstrates how different types of VTA neurons could collectively encode information about an ongoing series of actions.

### Cue and Reward Evoked Responses

In the first session, animals executed a single action to earn rewards (FR01). Similar to previous studies (Ljungberg et al., 1992; Schultz et al., 1993; Morris et al., 2006; Roesch et al., 2007), cue light onset at the start of the trial evoked homogenous, phasic population activation in VTA neurons. There was also a significant correlation between cue and reward delivery evoked responses. These observations are consistent with a convincing body of literature suggesting that dopamine neurons encode action value predicted by cues in the environment (Montague et al., 1996; Dayan and Abbott, 2001). On the other hand, when animals were required to execute multiple and random numbers of actions (random reinforcement schedules) to earn rewards we observed weakening of two critical neuronal responses by VTA neurons: (1)-loss of cue-evoked responses and (2) decorrelation of cue and reward evoked responses. This suggests that VTA neuronal responses to reward predictive cues are modulated by the contingency and/or contiguity between cues, action, and outcomes. When cues or discriminative stimuli are presented in FR01 trials, there is often a close temporal association between cue and reward delivery, as animals tend to execute actions within several seconds to earn rewards. As only one action is required in this setting, there may be a more accurate predictive relationship between cue onset and reward delivery. Thus, the unpredictable nature of a random reinforcement schedule may prevent the cue from evoking a reward prediction error. This finding is important because while previous studies have suggested that cue-evoke reward prediction errors can be used to guide action selection in a wide variety of settings, these data are drawn largely from Pavlovian and FR01-based tasks.

Cue offset in the current task is similar to a traditional Pavlovian conditioned stimulus, as offset was completely predictive of subsequent reward delivery with high temporal contiguity (0.5 sec between offset and delivery). Consistent with previous studies in which Pavlovian associations evoke VTA responses after associations are formed (Schultz, 1998), responses evoked by cue offset grew larger in successive sessions of the task. Our experiment was not designed to dissociate the effects of learning from the effects of different reinforcement schedules on information processing, and future experiments can directly address this topic. Taken together, these data suggest VTA neurons encode real-time information about ongoing actions in conjunction with information about cues with strong contiguity and contingency with reward delivery.

## Conclusions

Our date reveal a novel form of information processing by VTA neurons. The unique ensemble coding scheme that we observed allows heterogeneous groups of VTA neurons to provide a real-time account of ongoing behavior. This coding scheme may subserve the well-established role of dopamine in goal-directed actions, decision making, and the behavioral disorganization and amotivation associated with illnesses such as ADHD and schizophrenia.

## ACKNOWLEDGEMENTS

Funding was obtained from the NIMH MH048404 (BM), MH064537 (SK), MH064537 (REK), MH016804 (JW); NIDA DA031111 (JW), DA035050 (NWS), DA022762 (SK); RK Mellon Foundation (SK); Andrew Mellon Fellowship (JW). JW designed experiments, performed experiments, analyzed data, and wrote the manuscript. NWS designed experiments and edited the manuscript. SK analyzed the data. REK analyzed data and edited the manuscript. BM designed experiments and wrote the manuscript. The authors have no conflicts of interest related to this work.

---

**Fig. S1.**
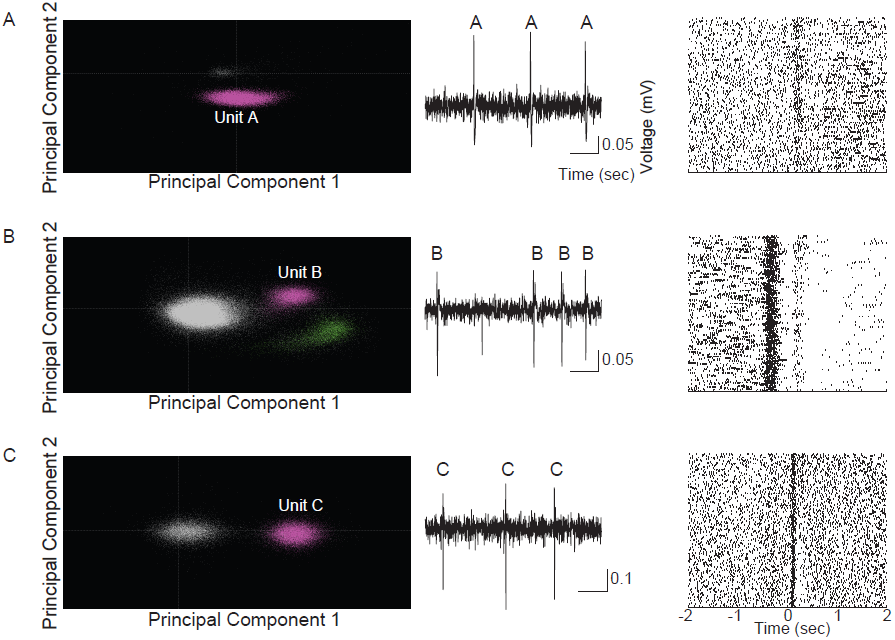
Representative examples of neuronal spike sorting, raw data and neuronal responses. (A) Spikes from unit A, a dopamine neuron, sorted according to the first 2 principal components of all threshold crossing waveforms (left). Purple points represent spikes that were assigned to unit A, and gray points represent noise that was not sorted into single unit spikes. A raw voltage trace (band pass filtered between 100 Hz and 7 KHz) corresponding to the same unit is depicted in the middle column. Examples of spikes belonging to unit A are notated in the trace. Unit A represents a typical cue-responsive unit. Raster plot depicts the unit’s response aligned to cue onset (right). Each dash represents a single spike, and each row represents a single trial (first trial in the top row). Note the increased spike density just after cue onset (time 0) across all trials. (B) Representative cue-offset responsive non-dopaminergic neuron. Data plotted with the same conventions as (A). In this example recording, two units were simultaneously recorded (left). Raster (right) depicts data during the time period between cue offset and reward delivery (-0.5 – 0s), with spikes aligned to the time of reward delivery. Note the consistent increase in spike density after cue offset and preceding outcome delivery. (C) Data from a representative non-dopaminergic, outcome delivery responsive neuron is plotted with similar conventions as (A). Raster (right) depicts neuronal activity aligned to the time of outcome delivery. Note the consistent delivery evoked response.

